# Improving the chromosome-level genome assembly of the Siamese fighting fish (*Betta splendens*) in a university Master’s course

**DOI:** 10.1101/2020.03.06.981332

**Authors:** Stefan Prost, Malte Petersen, Martin Grethlein, Sarah Joy Hahn, Nina Kuschik-Maczollek, Martyna Ewa Olesiuk, Jan-Olaf Reschke, Tamara Elke Schmey, Caroline Zimmer, Deepak K. Gupta, Tilman Schell, Raphael Coimbra, Jordi De Raad, Fritjof Lammers, Sven Winter, Axel Janke

## Abstract

**Background:** Ever decreasing costs along with advances in sequencing and library preparation technologies enable even small research groups to generate chromosome-level assemblies today. Here we report the generation of an improved chromosome-level assembly for the Siamese fighting fish (*Betta splendens*) that was carried out during a practical university Master’s course. The Siamese fighting fish is a popular aquarium fish and an emerging model species for research on aggressive behaviour. We updated the current genome assembly by generating a new long-read nanopore-based assembly with subsequent scaffolding to chromosome-level using previously published HiC data.

**Findings:** The use of nanopore-based long-read data sequenced on a MinION platform (Oxford Nanopore Technologies) allowed us to generate a baseline assembly of only 1,276 contigs with a contig N50 of 2.1 Mbp, and a total length of 441 Mbp. Scaffolding using previously published HiC data resulted in 109 scaffolds with a scaffold N50 of 20.7 Mbp. More than 99% of the assembly is comprised in 21 scaffolds. The assembly showed the presence of 95.8% complete BUSCO genes from the Actinopterygii dataset indicating a high quality of the assembly.

**Conclusion:** We present an improved full chromosome-level assembly of the Siamese fighting fish generated during a university Master’s course. The use of ~35× long-read nanopore data drastically improved the baseline assembly in terms of continuity. We show that relatively in-expensive high-throughput sequencing technologies such as the long-read MinION sequencing platform can be used in educational settings allowing the students to gain practical skills in modern genomics and generate high quality results that benefit downstream research projects.

## Introduction

The Siamese fighting fish, *Betta splendens*, is known for its eponymic aggressive behaviour between conspecific males. It was introduced into the international aquarium trade from the wild almost 130 years ago. The wildtype of *B. splendens* is endemic to Thailand and inhabits intact marshlands in shallow zones [1]. It is classified as “vulnerable” by the International Union for Conservation of Nature (IUCN) with decreasing population trends due to habitat destruction and pollution [1]. As a popular aquarium fish, it has been under strong artificial selection to produce several morphotypical variants as well as heightened aggressive behaviour. Numerous studies have focused on the psychological [2], behavioural [3], and ecological aspects [4] of this artificial selection. Genetic studies mostly investigated the genetic basis of the manifold of colours and fin shapes found in this species [5, 6].

Recently, [7] generated a chromosome-level *B. splendens* reference assembly. In order to do so, they first generated a baseline assembly using a combination of paired-end and mate pair libraries (sequenced on the Illumina HiSeq2000 platform), and then super-scaffolded the resulting assembly using a proximity-ligation-based HiC library (sequenced on the BGISEQ-500 platform). To further improve this assembly and to provide a solid basis for future analyses on this important fish model, we generated a more continuous baseline assembly using long-read data generated with the MinION sequencing device from Oxford Nanopore Technologies (ONT), and subsequently carried out scaffolding using the published HiC data from [7].

Data generation and genome assembly was carried out by students in the framework of a six-week Master’s course. This demonstrates the great potential of newly developed genome sequencing technologies for education. We hope that our study encourages academic institutions to offer hands-on genomics courses to students to gain first-hand experience in working with genomic data.

## Data Description

### DNA Extraction and Sequencing

We extracted high molecular weight DNA from muscle tissue of two individuals of aquarium-kept Siamese fighting fish using the protocol described in [8]. DNA quantity and fragment lengths were checked using the Genomic DNA ScreenTape^®^ (TapeStation Analysis Software A.02.01 SR1). We prepared four sequencing libraries using ONT’s Rapid (SQK-RAD004; three libraries) and 1D (SQK–LSK109; one library) sequencing kits. The resulting libraries were sequenced on the ONT MinION platform. All four sequencing runs yielded a total of ~21 Gbp of read data, with an average read length N50 of ~5.8 Kbp, ranging from 1.2 to 8.6 Kbp for the different sequencing runs (Supplementary Figure 1 and Supplementary Table 1).

### Genome Assembly and Scaffolding

We used Albacore v.2.3.3 (https://community.nanoporetech.com) for basecalling of the raw reads, which resulted in 18.7 Gbp of read data after removing reads with average quality scores below 7. In order to generate an overlap-layout graph for subsequent assembly, we first used Minimap2 v.2.14-r883[9] to carry out all-versus-all mapping using the default parameters for ONT data. Subsequently, we used Miniasm v.0.3-r179 [10] to generate the assembly graph and call the consensus sequence. The resulting genome showed a size of 441 Mbp with 1,276 contigs and a N50 of 2.1 Mbp. For consensus polishing, we first aligned the nanopore reads back to our assembly using Minimap2 and performed the error correction using Racon v.1.3.1 [11]. This step was repeated twice. Next, to further improve the resulting consensus quality, we carried out error correction using previously published Illumina paired-end short-read data (accession no. SRR6251365;[7]). For that, we first used Cutadapt v.1.18 [12] to remove adapter sequences as well as low-quality ends from the reads. We then mapped the paired-end (SRR6251365) and mate pair (SRR6251353) data onto the genome assembly using BWA-MEM v.0.7.17-r1188 [13] and sorted the resulting mapping file using SAMtools v.1.9 [14]. Lastly, we carried out the polishing for three rounds using Pilon v.1.23 [15].

In order to achieve chromosome-level for our long-read based assembly, we removed all contigs matching to the mitochondrial genome, and subsequently mapped the previously published HiC reads (accession no. SRR6251367;[7]) onto the genome using BWA-MEM. Next, we scaffolded the assembly using the HiC reads with ALLHiC v.0.9.8 [16]. This resulted in 109 scaffolds with a scaffold N50 of 20.7 Mbp (Table 1). Over 99% of the assembly size was placed into 21 chromosomes. A contact map of the resulting assembly can be seen in Figure 1A. The genome assembly and all read data generated during this project are accessible on GenBank (Bioproject PRJNA592275).

**Table 1:** Genome continuity statistics for the HiC scaffolded genome calculated with QUAST.

|  | HiC scaffolded assembly |
| --- | --- |
| Number of contigs/scaffolds | 109 |
| Largest scaffold | 34.1Mb |
| Scaffold N50 | 20.7 Mb |
| Scaffold L50 | 9 |
| Assembly size | 441.2 Mb |
| GC (%) | 45.1 |

**Figure 1:**
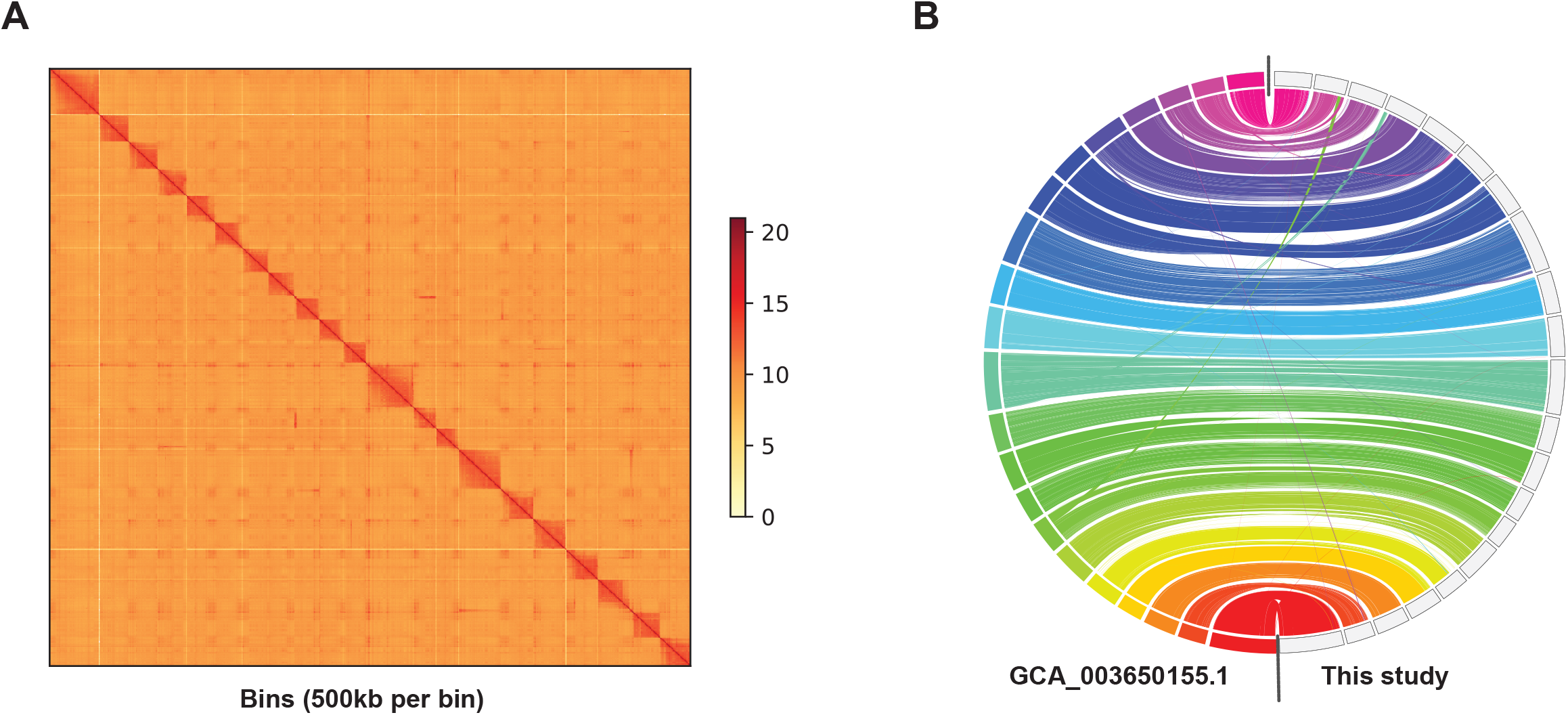
(A) HiC contact map of the 21 chromosome-level scaffolds, and the shorter unplaced scaffolds. As can be seen in the plot, the assembly only shows small amounts of trans-chromosomal interactions. (B) Whole genome synteny between the chromosome-level assembly of [7] (on the left) and our chromosome-level assembly (on the right). The lines indicate aligned regions between the two assemblies.

### Genome Quality Assessment

We obtained genome continuity statistics (Table 1) with QUAST v5.0.2 [17] and assessed assembly completeness using BUSCO v.3.0.2 [18] with the Actinopterygii gene set (busco.ezlab.org). We then mapped the Illumina HiSeq2000 reads from the 5 Kbp insert-size mate pair library (accession no. SRR625353;[7]) to the assembly and investigated the distribution of insert sizes for the library (Supplementary Figure 2). We mapped the data using BWA-MEM, sorted the alignment files with SAMtools, marked duplicates using Picard 2.20.7 (https://broadinstitute.github.io/picard/), and then created a histogram based on the statistics obtained from GATK 4.1.4.1 (https://gatk.broadinstitute.org; CollectInsertSizeMetrics option). We observed a much higher rate of read pairs mapping with the right insert size in both our polished nanopore baseline assembly and our final chromosome-level assembly compared to the chromosome-level assembly of [7]. Investigating synteny changes between the two chromosome-level assemblies with JupiterPlot [19], we found a strong overall agreement with some differences especially towards the ends of the scaffolds (Figure 1B). We further investigated the amount of contaminated contigs in our assembly using Blobtools 1.1.1 [20]. The analysis showed no signs of contamination, since 99.99% of the assembly were taxonomically assigned as Chordata and the majority of the scaffolds and contigs showed highly similar coverage and GC contents (Supplementary Figure 3). We found very narrow peaks for the distributions of coverage and GC content in the assembly, with only a few short contigs showing slightly lower GC content than the majority of the genomic contigs/scaffolds.

### Transcriptome Assembly and Quality

In order to assemble the transcriptome of *B. splendens* for subsequent use in gene annotation, we downloaded seven previously published RNA-seq libraries from NCBI (accession no. SRR6251368–SRR6251375). We assembled the transcriptomes *de novo* using Oases v.0.2.09 [21]. The completeness of the transcriptome assembly was assessed with BUSCO, using the Actinopterygii gene set, revealing 93.4% complete, 4% fragmented, and 2.6% of missing BUSCO’s.

### Genome annotation

#### Repeat annotation

In order to annotate repeats in our assembly we first created a custom *de novo* repeat library using RepeatModeler v.1.0.11 (www.repeatmasker.org/RepeatModeler/) and then combined this library with the curated *Danio rerio* repeat dataset from Dfam 3.0. Repeats in the genome were then annotated and masked using RepeatMasker open-4.0.7 (www.repeatmasker.org/RepeatMasker/). The results are provided in Table 2.

**Table 2:** Repeat content of the HiC scaffolded assembly.

| Type of element | Number of elements | Length | Percentage of assembly |
| --- | --- | --- | --- |
| SINEs | 12,939 | 2,732,859 | 0.62% |
| LINEs | 51,088 | 31,431,107 | 7.13% |
| LTR elements | 24,596 | 21,574,199 | 4.89% |
| DNA transposons | 54,813 | 14,902,018 | 3.37% |
| Unclassified | 74,783 | 23,656,821 | 5.36% |
| Small RNA | 6,578 | 1,201,687 | 0.27% |
| Satellites | 2,397 | 1,216,117 | 0.28% |
| Simple repeats | 385,394 | 24,039,213 | 5.44% |
| Low complexity | 35,821 | 2,031,477 | 0.46% |
|  |  | **Total:** | 27.82% |

#### Gene annotation

Gene annotation was performed using MAKER2 v.2.31.10 [22] in several steps. First, we carried out evidencebased annotation using proteins obtained from [7] (available at gigadb.org/dataset/100433) and our aforementioned *de novo* assembled transcriptomes. We then trained the *ab initio* gene predictor SNAP v.2006-07-28 [23] using MAKER2 results over two rounds. Additionally, we used the two *ab initio* gene predictors Augustus v.3.3 [24] and Genemark v.4.38 [25]. This resulted in 21,535 annotated transcripts, which is slightly lower than the 23,981 gene models generated by [7].

We applied BUSCO and DOGMA 3.4 [26] to asses the completeness of the annotated proteome. Within BUSCO the Actipterygii set yielded 81.5% (n = 4,584) complete core orthologs and within DOGMA 83.88% (n = 8,113) of the vertebrate sets conserved domain arrangements (CDAs). About 90% of all gene models showed Annotation Edit Distance (AED)scores < 0.5, indicated a high quality (Supplementary Figure 4). An AED score of 0 indicates perfect agreement of the evidence and the gene prediction, and a score of 1 indicates that the gene prediction is not supported by any evidence [27].

## Educational aspect of the assembly generation

The MinION’s potential as an effective teaching tool was recognized early on and has been used in classroom settings [28-30] as well as in the field [31, 32]. Here we show that inexpensive nanopore-based sequencing along with memory and run-time efficient genome assembly tools offer great potential to generate high quality chromosome-level assemblies, even of more complex vertebrate genomes, as part of university courses. Scientific topics like high-throughput sequencing, the bioinformatics of genome assembly and genome evolution can thereby be taught in a highly applied and engaging way. Furthermore, modern technologies do not only offer the chance for invaluable training using state-of-the-art methods, but also allow students to publish results early on in their career. The ever-decreasing sequencing costs should enable universities, even in low-income areas and countries, to train their students in modern genomics and bioinformatics.

## Authors’ Contributions

SP and AJ designed the study. SW, MG, SJH, NKM, MEO, JOR, TES and CZ carried out the laboratory procedures and the sequencing. SP, MP, MG, SJH, NKM, MEO, JOR, TES, CZ, DG, TS, RC, MNJ, JDR, FL and SW carried out the bioinformatic processing and analyses. All authors contributed to writing this paper.

## Acknowledgments

We thank Damian Baranski for help with the DNA extraction, and the LOEWE-Centre for Translational Biodiversity Genomics and the Goethe University, Frankfurt for providing the financial and practical resources required to perform this study and run the course. The present study is a result of the Centre for Translational Biodiversity Genomics (LOEWE-TBG) and was supported through the programme “LOEWE – Landes-Offensive zur Entwicklung Wissenschaftlich-ökonomischer Exzellenz” of Hesse’s Ministry of Higher Education, Research, and the Arts.

## Supplementary Figure Legends

**Supplementary Figure 1: Comparison of the data output and read lengths between all four MinION sequencing runs.** Run 1: Oxford Nanopore Technologies (ONT) 1D sequencing kit (SQK–LSK109) and Run 2-4: ONT Rapid sequencing kit (SQK-RAD004). A) Read quality scores of the four different runs, B) bog-transformed read lengths, C) numbers of reads, and D) the total amount of sequencing data generated.

**Supplementary Figure 2: Insert sizes of the 5 Kbp mate pair read library of [7], mapped against the chromosome-level assembly of [7] (GCA_003650155.1; top panel), our nanopore-based baseline assembly (middle panel) and our chromosome-level assembly (bottom panel).**

**Supplementary Figure 3: Blobtools plot showing the taxonomic assignments (blue colour for Chordata, and gray for “no hits”) of the different scaffolds, and scaffold-wide coverage and GC contents.** The scaffolds were blasted against the NCBI nucleotide database.

**Supplementary Figure 4. Distribution of Annotation Edit Distance (AED) scores.** About 90% of all gene models show AED scores of < 0.5 indicating a high quality of our gene models.

**Supplementary Tables**

**Supplementary Table 1:** Statistics of the four MinION sequencing runs.

|  | Run 1 | Run 2 | Run 3 | Run 4 |
| --- | --- | --- | --- | --- |
| **Library Type** | SQK-LSK109 | SQK-RAD004 | SQK-RAD004 | SQK-RAD004 |
| **Mean read length** | 793 | 2,845 | 4,049 | 3,505 |
| **Mean read quality:** | 7.5 | 8.2 | 8.9 | 8.9 |
| **Number of reads:** | 5,619,880 | 937,354 | 1,253,758 | 2,526,439 |
| **Read length N50:** | 1,158 | 6,654 | 8,556 | 6,919 |
| **Total bases:** | 4,454,832,013 | 2,666,680,634 | 5,076,315,340 | 8,855,545,432 |

