## Supplementary figures and images for "Improving the chromosome-level genome assembly of the Siamese fighting fish (*Betta splendens*) in a university Master’s course"

### Supplementary Figure 1

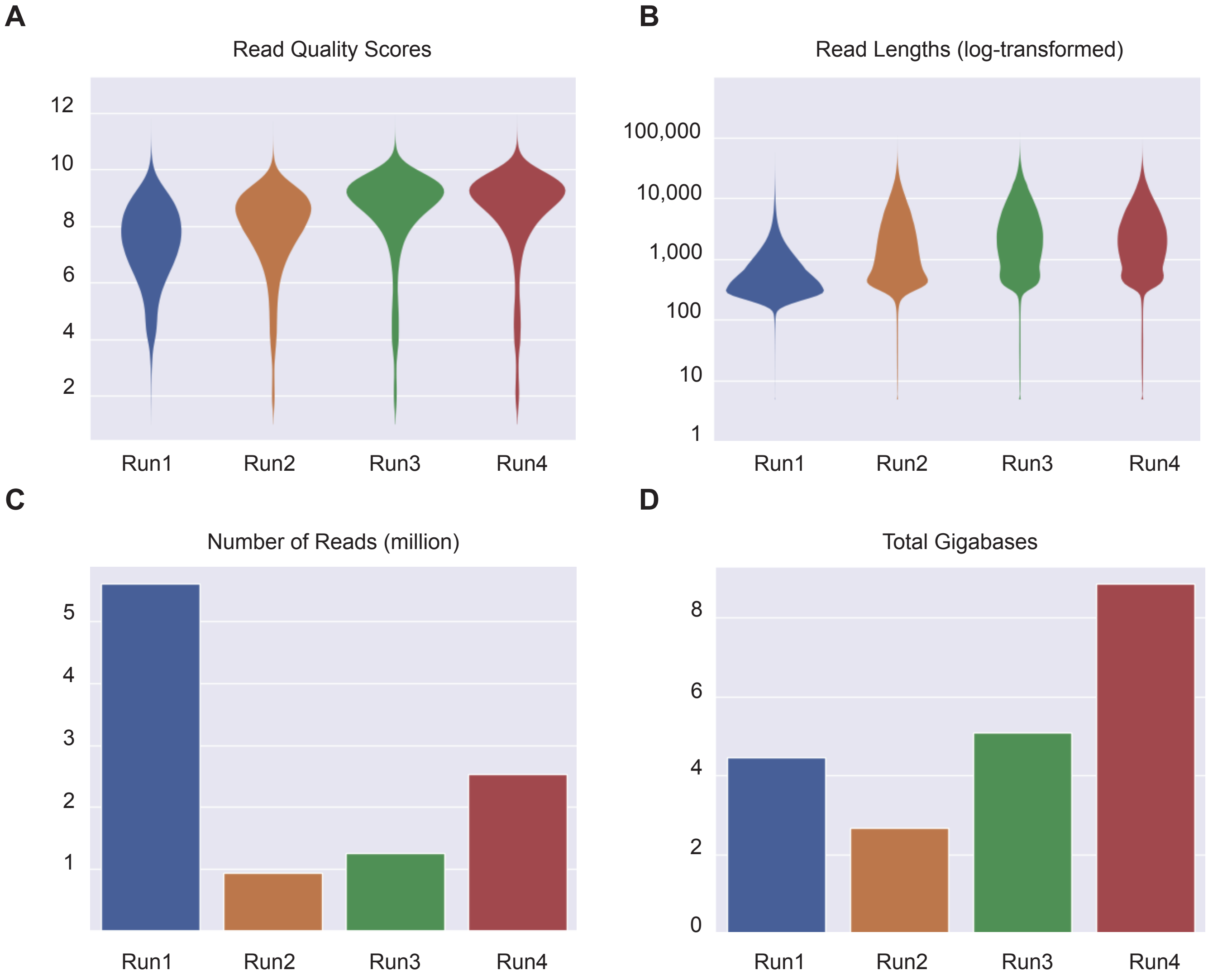

### Supplementary Figure 2

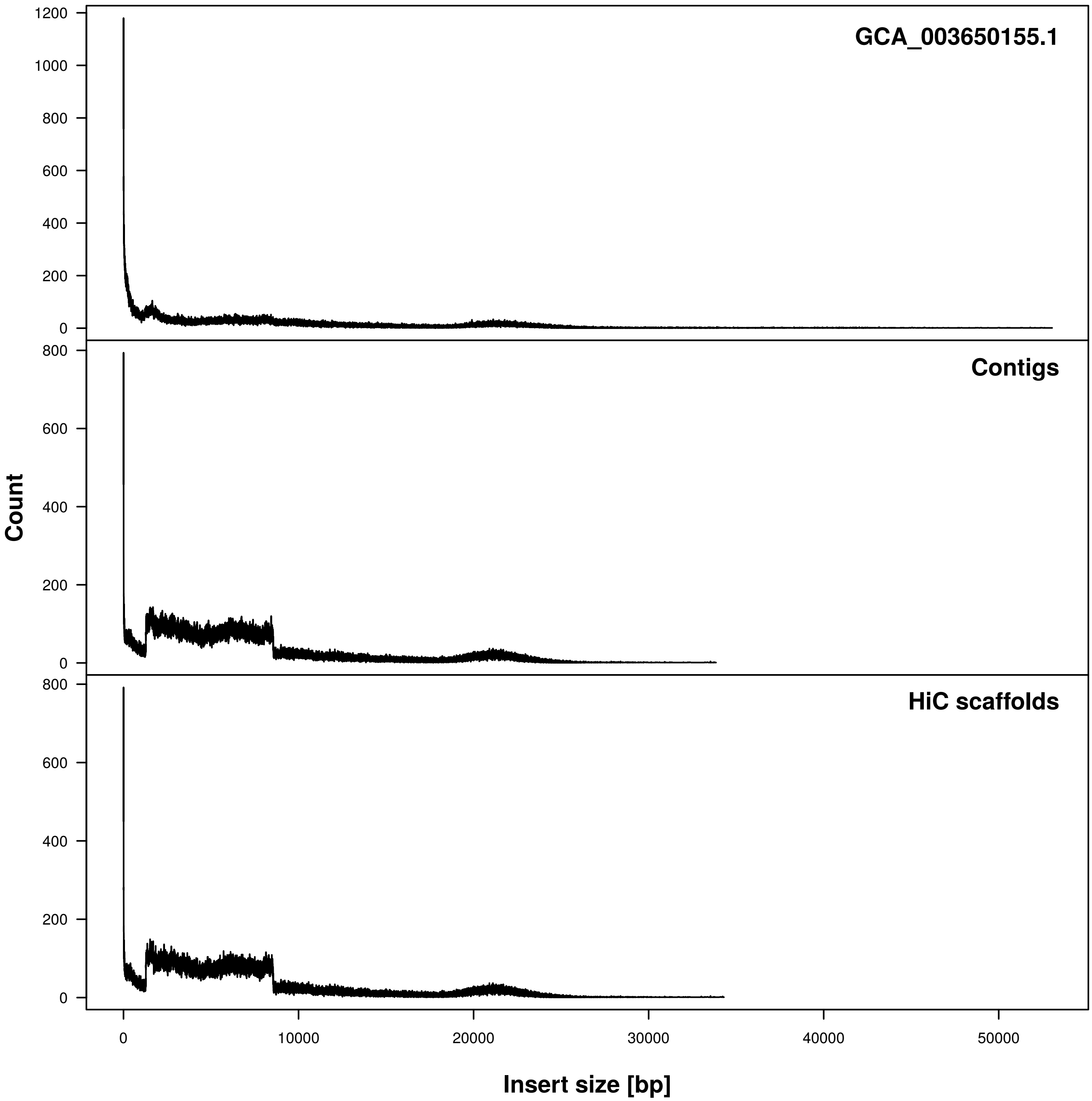

### Supplementary Figure 3

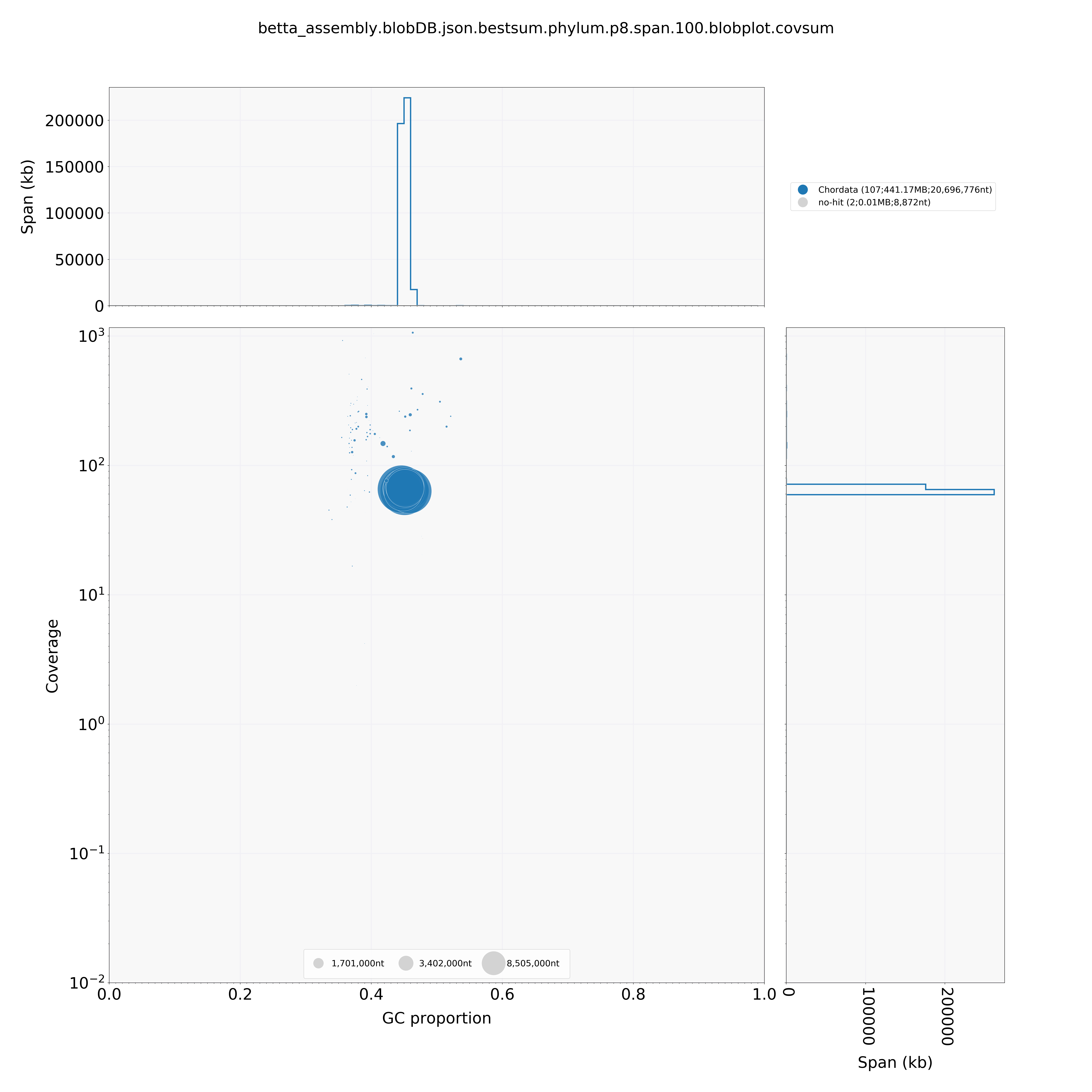

### Supplementary Figure 4

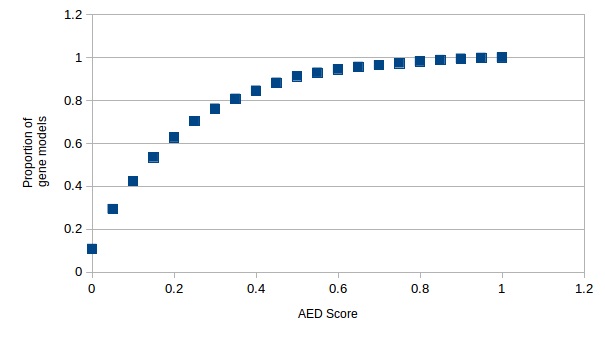
